# Aiptasia oral regeneration is host-controlled but supported by symbiont-derived photosynthates

**DOI:** 10.1101/2025.03.10.642338

**Authors:** Jun B. Cai, Samuel Bedgood, Virginia M. Weis, Lucía Pita

**Author notes:** Corresponding Author: Lucía Pita, Institute of Marine Research, Spanish National Research Council (IIM-CSIC), Eduardo Cabello 6, Vigo, Pontevedra, 36208, Spain. Division of Natural Sciences, New College of Florida, Sarasota, FL, United States of America.

## Abstract

The study of oral regeneration in the sea anemone *Exaiptasia diaphana* (commonly called Aiptasia) - a prominent model for the study of coral-algal symbiosis - offers the unique opportunity to investigate the role of symbionts in regeneration. Algal symbionts have the potential to affect healing and regeneration by supplementing host nutrition and/or modulating host immune responses. Here we revisited the descriptions of Aiptasia oral regeneration from the 1970s with modern imaging techniques. Upon oral amputation, the anemones progressed through six distinct and repeatable stages of wound healing and regeneration, a process that was completed within a week. We followed regeneration in symbiotic anemones (i.e., associated with native dinoflagellate symbionts) in comparison to aposymbiotic anemones (i.e., lacking algae), and symbiotic anemones kept in the dark (i.e., blocking photosynthesis). In most symbiotic anemones under normal light, tentacle buds appeared within 32 hours post-amputation, whereas in aposymbiotic anemones and symbiotic anemones kept in the dark buds appeared 12 hours later. This pattern suggests that the contribution of symbiont-derived photosynthates to host nutrition shortened regeneration time. Our study provides the basis for further research on the underlying molecular processes in Aiptasia oral regeneration, for comparative studies of different symbiotic cnidarians and for investigating the relative role of host and symbionts in different developmental processes such as whole-body morphogenesis during asexual reproduction.

## Introduction

Wound healing and regeneration are key restoration processes in all animals, although regenerative capacity varies greatly across taxa (Bely & Nyberg, 2010; Slack, 2017). Among animals capable of whole-body regeneration are several cnidarians, including the hydroids *Hydra, Hydractinia* and *Tubularia* (Bosch, 2007; Barth, 1944; Bradshaw, Thompson & Frank, 2015; Murad et al., 2021) and the anthozoan sea anemone *Nematostella* (Passamaneck & Martindale, 2012; Amiel et al., 2015; Schaffer et al., 2016; Warner et al., 2018; DuBuc, Traylor-Knowles & Martindale, 2014). Besides some commonalities, such as Wnt-signaling-mediated patterning (Bosch, 2007; DuBuc, Traylor-Knowles & Martindale, 2014), how these organisms achieve whole-body regeneration also varies, including the cell types involved and the relevance and timing of cell proliferation (Bosch, 2007; Passamaneck & Martindale, 2012). The sea anemone *Exaiptasia diaphana* (commonly called Aiptasia) also displays high regenerative capacities (Singer & Palmer, 1969a,b; van der Burg et al., 2020; van der Burg & Prentis, 2021), and it is a prominent model system for the study of coral-algal symbiosis (Weis et al., 2008; Rädecker et al., 2018; Weis, 2019). To date, there has been no attention given to the role of symbiotic state in the host animal regeneration process.

Processes in which homeostasis is disrupted, such as injury, offer the opportunity to assess the influence of the microbiota on animal health (Shavandi et al., 2020). A breach of animal physical barriers will induce an inflammatory response (LeBert & Huttenlocher, 2014; Wenger et al., 2014; Wu et al., 2022), modulate cell division and cell death (Ricci & Srivastava, 2018; Song et al., 2020), and require metabolic investment (Henry & Hart, 2005; Starostová, Gvoždík & Kratochvíl, 2017), (see also the review by Rennolds & Bely, 2023 and references therein). All three of these processes can be modulated by symbiotic microbes. For example, the microbiota induces or silences inflammatory responses (reviewed in Belkaid and Hand 2014), endosymbionts and their host cells coordinate cell division (Tivey, Parkinson & Weis, 2020), and one key role of the microbiome is contributing to host nutrition, as shown in multiple systems (Luan et al., 2015; Pita et al., 2018; Fontaine & Kohl, 2020).

The study of whole-body regeneration in Aiptasia and the potential role of symbionts in the process goes back to pioneering papers by Singer and collaborators. Upon oral disc amputation, tentacle buds appear 3 days post-amputation (Singer & Palmer, 1969a), preceded by proliferation of epidermal cells –but with little-to-no proliferation of gastrodermal cells (Singer, 1971). Although oral regeneration is a metabolically costly process (Singer & Palmer, 1969b), the authors did not detect remarkable differences between the regeneration rate of anemones with no or few symbiotic algal cells *vs* anemones fully colonized by symbionts (Singer & Palmer, 1969a). During pedal lacerate development, a form of asexual reproduction in anemones that resembles whole-body regeneration, aposymbiotic anemones develop tentacles earlier than symbiotic ones, but then anemone growth rate, as measured by tentacle number, in symbiotic anemones overtakes aposymbiotic anemones (Presnell, Wirsching & Weis, 2022). A working hypothesis to explain this is that symbiont-induced immune suppression to promote tolerance (Weis 2019) interferes with wound-induced inflammation at the early stages of regeneration. In the later stages of development, the presence of symbionts promotes growth rate via increased nutrition from algal photosynthesis and/or growth signals from the symbionts to the host (Presnell, Wirsching & Weis, 2022; Tivey, Parkinson & Weis, 2020). Here we revisited the work on Aiptasia oral disc regeneration by Singer and collaborators to test the contribution of symbionts to the process. We characterized the different stages of healing and oral regeneration in Aiptasia. We then evaluated if symbiosis with the native photosynthetic dinoflagellate *Breviolum minutum* (Family Symbiodinaceae) influenced the timing of healing, formation of tentacle buds, and tentacle growth during oral regeneration.

## Materials & Methods

### Aiptasia maintenance and oral disc amputation

Experimental anemones were generated from stocks of the clonal H2 strain containing native dinoflagellate *Breviolum minutum*, originally collected from a single individual from Coconut Island, Hawaii (Thornhill et al., 2013; Xiang et al., 2013). Symbiotic anemones were maintained on a 12h:12h light:dark photoperiod (approximately 20 µmol quanta m-2 s-1) at 25°C. Aposymbiotic anemones were maintained in the dark at 25°C. Anemone stocks were fed 3 times per week with live *Artemia* nauplii and filtered artificial seawater (FSW; Instant Ocean) was replaced several hours after feedings. Experimental anemones were starved one week prior to the beginning of experiments and not fed for the duration of the experiments.

Aposymbiotic anemones were generated ∼1 month before all experiments following a modified menthol-bleaching method from Matthews et al., 2016. In short, symbiotic anemones were incubated in a 0.58 mM solution of FSW and menthol (Sigma Aldrich) dissolved in ethanol for 4h daily for 4 days. This was repeated weekly until anemones were symbiont-free as assessed by a Zeiss Axio Observer A1 fluorescence microscope. Aposymbiotic status was confirmed before the start of experiments.

Four to five days before the beginning of each experiment, small anemones (>2.5-5 mm pedal discs) with 12-15 tentacles were placed in 6-well plates containing a glass coverslip. Anemones attached to the coverslip were easiest to manipulate for oral disc amputation and visualization; thus, unattached anemones were removed. We relaxed the anemones for 20 minutes in a 1:1 solution of 7% MgCl_2_ in FSW. Oral disc amputation was performed with a scalpel blade #15C, by transverse dissection at the middle of the body column. The relaxing solution was then removed and replaced with FSW.

### Morphological changes during wound healing and oral regeneration

During a 1-week period, 59 anemones were observed and imaged using a Canon EOS 60D digital camera under a Leica M165 FC stereo microscope, and the different stages of healing/regeneration were established based on phenotypic changes, such as presence of acontia, shape, presence of buds, elongation of tentacles. Anemones were checked at the following time points: 3 hours post-amputation (hpa), 5/6, 8, 24, 32, 48, 56, 96, 120, and 168 hpa. In a pilot experiment, we also recorded a time-lapse video of the first 4 hpa of an individual. Pictures were taken every 20s and then converted into a video at 20 frames per second.

### Oral regeneration under the microscope

At different stages of healing/regeneration, samples were fixed and processed for fluorescence staining of host cell nuclei (0.2 µg/mL Hoechst, Invitrogen) and actin filaments (0.0066 µM Alexa Fluor 488-phalloidin, Invitrogen) as described in Presnell, Wirsching & Weis, (2022). In short, animals were relaxed in 7% MgCl_2_ and then fixed in 3.7% formaldehyde/0.2% glutaraldehyde/1X PBS for 1 min at room temperature, then fixed in 3.7% formaldehyde/1X PBS and incubated overnight at 4 °C. After several washes in PBS with 0.2% Triton X-100, samples were incubated in this solution with 0.5% bovine serum albumin overnight. Samples were incubated in the dye for 8h, then washed in PBS several times before going into a series of washes of increasing concentrations of glycerol in PBS: 50% glycerol overnight at 4ºC, 70% glycerol for 15 min at room temperature, and 87% glycerol overnight at 4ºC. Samples were incubated and mounted in 87% glycerol. Images were acquired on an LSM780 NLO Confocal microscope (Zeiss) at the Center for Quantitative Life Sciences, Oregon State University, Corvallis, USA. Hoechst, phalloidin, and chlorophyll a (symbiont autofluorescence) Z-stack images were acquired on separate channels with a 405 nm Diode, an Argon, and 633 nm HeNe laser, respectively. Pinholes for each channel were set at 1.49 AU, 1.19 AU, and 0.99 AU, respectively. Images were processed in ZEN blue for pseudocoloring, without further processing.

In addition, we tested the effect of hydroxyurea (HU, Thermo-Fisher Scientific #A10831.03, Waltham, MA), an inhibitor of DNA replication, arresting cells in S-phase of their cycle on symbiotic anemones. We performed a dosage test with different concentrations of HU on control Aiptasia (i.e. intact polyps) over a one-week period, which included 5, 10, and 20mM HU concentrations. 10mM HU halted almost all cell division (EdU labeling described below), and 20mM HU killed the polyps (**Fig. S1**). 15mM showed inhibition of cellular proliferation without lethal effects. Prior to each experiment, HU was made fresh at 15 mM by diluting in FSW. HU treatment was performed on experimental anemones (n=46) after 6/8 hpa. The inhibitor solution was exchanged every 12h with freshly-diluted inhibitor. Control anemones (n=41) received FSW changes every 12h. Based on the different stages of healing/regeneration defined above, we monitored the progress of oral regeneration in each polyp. At the end of the experiment, anemones were incubated in 10µM 5-ethynyl-2’-deoxyuridine (EdU) Alexa Fluor 555 (Click-iT EdU Alexa Fluor 555 imaging kit; Life Technologies, Eugene, OR, USA) in FSW (with or without HU) for 12h to visualize host cell proliferation. EdU (-) controls were also included. Polyps were then processed as in Tivey, Parkinson & Weis, 2020. In short, polyps were relaxed and fixed in 1X PBS+ 4% paraformaldehyde, blocked and permeabilized, and then processed with Click-iT EdU reaction kit. All host nuclei were labeled with Hoechst 3342 (Invitrogen) as detailed above. Finally, polyps were washed three times in 1X PBS and mounted on slides for confocal microscopy (glycerol 87% in PBS). Confocal images were acquired as described above. EdU-labeled proliferative cells were detected with a 561 nm laser with a pinhole set at 1.01 AU. Symbiont autofluorescence and host nuclei labeled with Hoechst were detected as described above. An EdU negative control served to check for background fluorescence. Confocal Z-stack images were acquired.

### Healing and oral regeneration in symbiotic *vs* aposymbiotic anemones

We amputated the oral discs and recorded the different stages of regeneration in three different treatments: aposymbiotic anemones (n = 56), symbiotic anemones (n = 47), and symbiotic anemones kept in the dark by wrapping the plates in aluminum foil (to block photosynthesis and thereby eliminate the contribution of symbionts to host nutrition), (n = 24). We performed two consecutive experiments (i.e., 1 week apart) with these conditions to reach the total number of replicates per treatment indicated above. We recorded the stage of regeneration under a Leica M165 FC stereo microscope, as described above. Anemones in dark conditions were briefly exposed to light when inspected under the stereo microscope. When the position of the anemone did not allow us to clearly assess the stage of regeneration, the data were recorded as not available (NA), **Data S1**.

### Statistical analysis for comparison of regeneration under different treatments

We tested if the progress of regeneration in an individual was influenced by treatment by ways of generalized linear models in R 4.2. 0 (R Team Core 2019): *glm(Stage ∼ Treatment *hpa + (1*|*Individual), family = poisson*). We included the random effect of individual (1|Individual) because we have repeated measures. Statistical significance (p-value <0.05) was determined by Chi-square test. We also compared the effect of treatment on the onset of each stage separately (Y = hpa, x = treatment). We first tested homoscedasticity by Levene’s test. In case of homoscedasticity (Levene’s test non-significant), we used Kruskal-Wallis test. In case of heteroscedasticity (Levene’s test p-value < 0.05), we applied a non-binomial modified glm (glm.nb) as implemented in the package MASS (Venables & Ripley, 2002) and statistical significance (p-value <0.05) was determined by Chi-square test. Post-hoc comparisons were calculated with Tukey test as implemented in the package emmeans (Lenth RV 2022). Data were visualized with the r package ggplot2 (Wickham 2016).

## Results

### Overview of oral regeneration in Aiptasia

We identified different stages of oral regeneration in Aiptasia, based on our observations during the first week after amputation (**Fig. 1)**. Upon oral disc amputation, anemones released acontia, filaments typically loaded with nematocytes that act as a defensive mechanism (stage 1). The wound is totally open, acontia and mesenteric filaments may also be observed at the wound. By 3h post-amputation (hpa), most anemones stopped releasing acontia and the wounded edges began to roll inward (stage 2, **Fig. 1A**) by contraction of the columnar circular musculature (**Fig. S2**). Although the wound seemed open, hydrostatic pressure was at least partially restored, as evidenced by animals being able to move across the substrate using their hydrostatic skeleton, (**Video S1**, 4 hpa). As the bending of the wounded edges progressed, the open wound closed and the anemones adopted an onion shape (stage 3, **Fig. 1A**). By 24 hpa, >90% of anemones had reached this stage (**Fig. 1B**). Note that we could not establish the exact time of wound healing. We considered polyp movement as an indication of re-established hydrostatic pressure. We observed that the polyps often changed position during the first hours post amputation, and until the “onion shape” was adopted. Therefore, we expect wound healing occurred during that time window, even when movement of the polyps was not evident. Afterward, the body column extended longitudinally and the first tentacle buds appeared (stage 4, **Fig. 1A**). It took between 24 h and 96 h for the appearance of tentacle buds, but most anemones reached stage 4 within 32-48 hpa (**Fig. 1B**). Tentacle buds appeared as a swelling of the epidermis, with retractor muscles still absent (**Fig. S2**). Treatment with HU arrested oral regeneration at stage 3 and did not elongate their body column or develop tentacle buds **(Fig. S3**). At stage 5, the first tentacles were fully emerged (stage 5, **Fig. 1A**). In some cases, a primary tentacle appeared first, but often several tentacles appeared at once. By this stage, the oral structure was complete, the musculature of the tentacle was formed, and the tentacles were fully colonized by algal symbionts (**Fig. S2**). Note that the regenerated mouth was not always visible, depending on the position of the polyp, so we assigned anemones to stage 5 according to the presence of tentacles. Finally, we defined stage 6 as the stage in which anemones showed ≥ 12 tentacles. Most anemones completed the process within 1 week (**Fig. 1B**). Additional representative images for each stage are provided in **Fig. S4**.

**Fig. 1.**
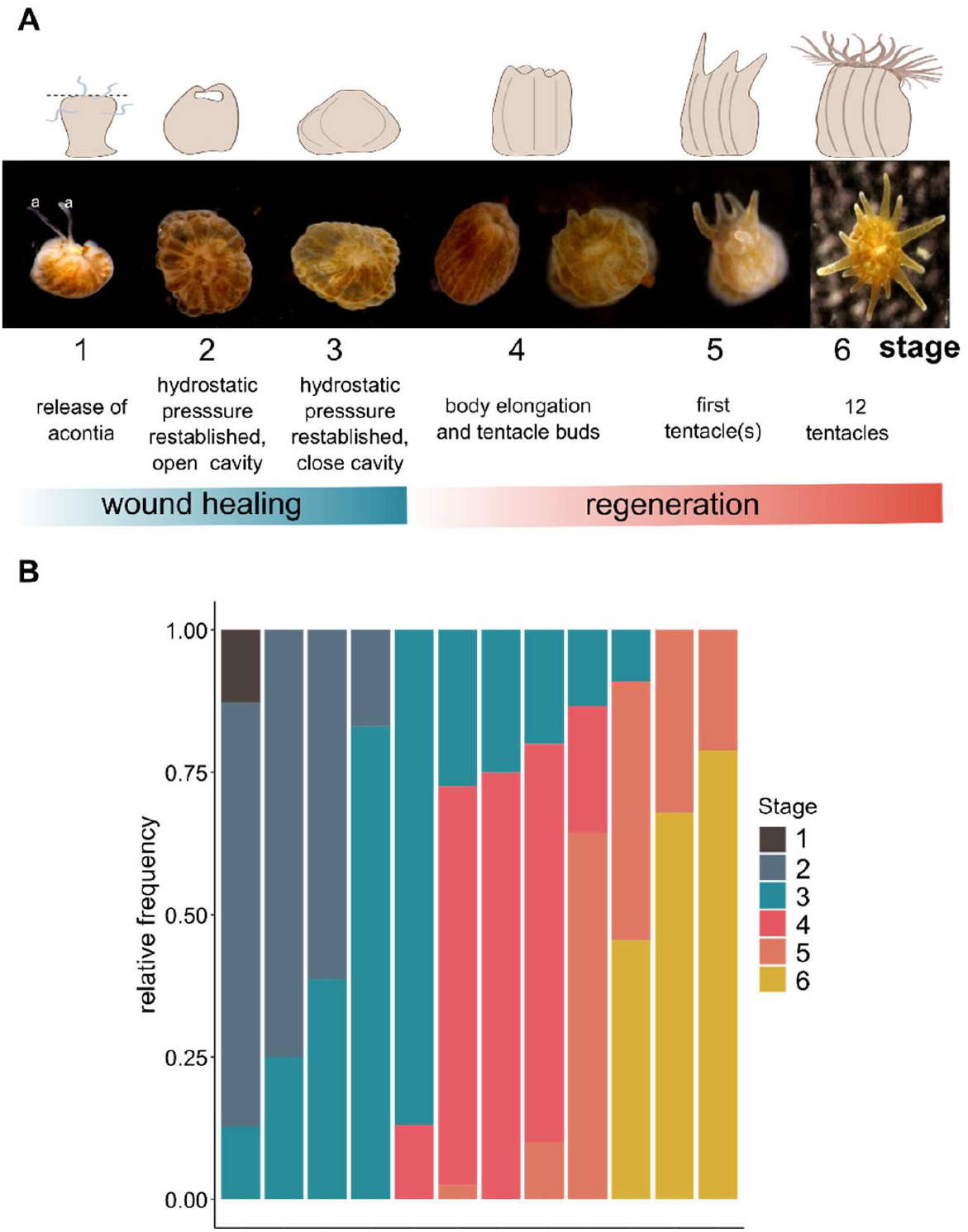
Overview of stages of Aiptasia oral regeneration. **(A)** Representative images of the regeneration stages we defined. Stage 1: the wound was open, anemones released acontia (a). Stage 2: the wound appeared open, yet the epidermis rolled inward. We counted anemones as reaching stage 2 at 3 hpa, as long as there were no acontia and the wounded edges were rolled in. Stage 3 (the onion shape): the restoration of hydrostatic pressure was clear; anemones appeared rounded and the wound was hardly visible. Stage 4: the anemone column elongated, and the first tentacle buds appeared. Stage 5: the first tentacles were evident. Stage 6: ≥ 12 tentacles. Aiptasia drawings were created with BioRender.com. **(B)** Progression of regeneration over time (hpa). Relative frequency of anemones at each regeneration stage at each time point (number of anemones at a particular stage divided by the total number of counted anemones at the given time point).

### Comparison of oral regeneration in aposymbiotic *vs* symbiotic anemones

We compared progression of regeneration in symbiotic anemones kept in a normal light regime (12h light:12h dark conditions), aposymbiotic anemones (without symbiotic algae, **Fig. S5**) in a normal light regime, and symbiotic anemones kept in the dark (24h dark conditions). Dark conditions prevent the photosynthetic activity of the symbionts; thus, there is no photosynthate supplied to the anemone from the algae. We observed no effect of treatment in progression of regeneration (glm (Chi) test, 1 degree of freedom, P = 0.9909). However, when we tested the time to reach each stage separately, we detected that stage 4 (i.e., appearance of tentacle buds) occurred significantly sooner in symbiotic anemones in a normal light regime compared to the dark treatment and the aposymbiotic anemones (glm, 2 degrees of freedom, P = 0.0002), (**Fig. 2, Table S1**).

**Fig. 2.**
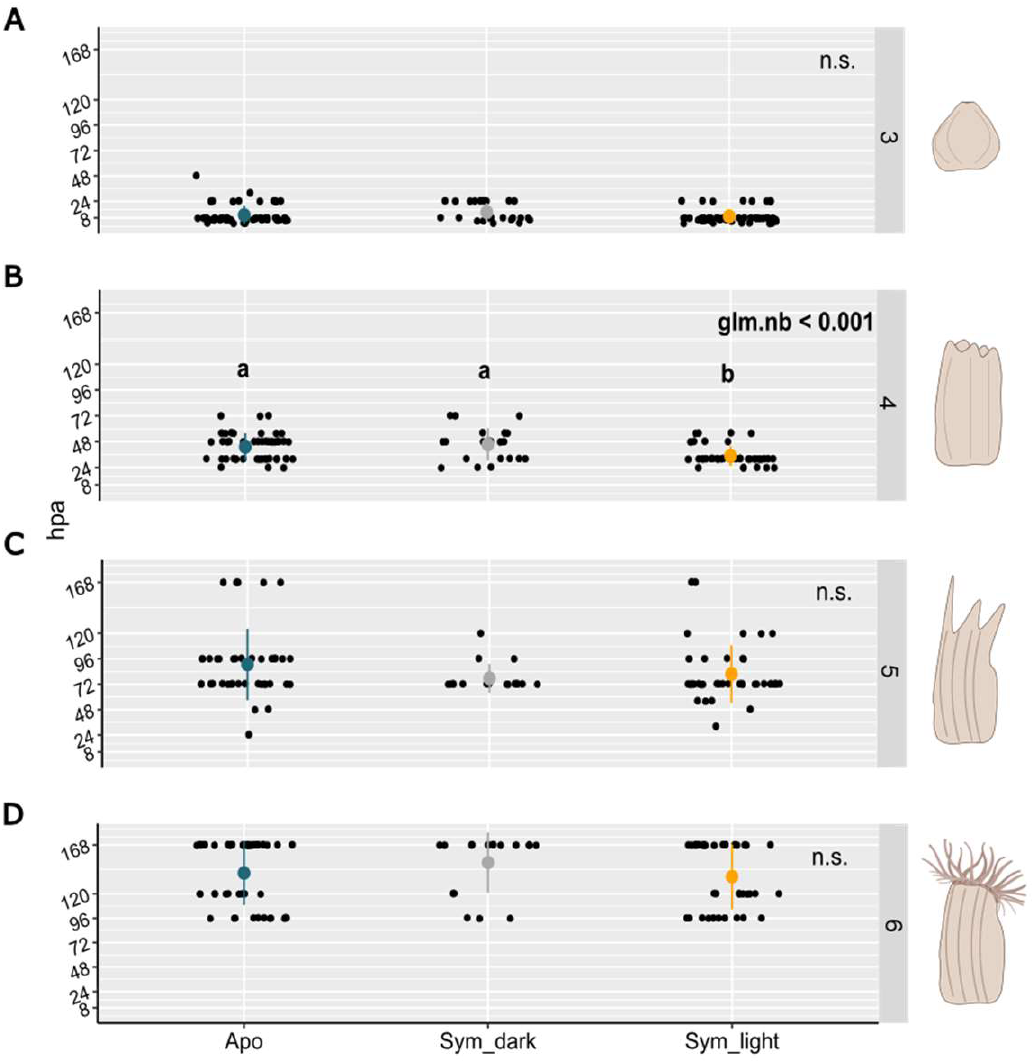
Progression of oral regeneration (time to stage) in different symbiotic states and light conditions. Time to reach **(A)** stage 3, **(B)** stage 4, **(C)** stage 5, and **(D)** stage 6. Symbiotic anemones in a normal light regime took similar time to reach stages 3, 5, and 6 as the other treatments; but the former reached stage 4 significantly faster. Mean ± standard deviation is shown for each treatment group in color. For stage 4 **(B)**, non-binomial glm test was significant, p(Chi) < 0.00. Letters represent grouping of treatments according to Tukey’s test results: similar treatments are designated with the same letter. hpa: hour post-amputation. Apo: aposymbiotic anemones; Sym_dark: symbiotic anemones growing in the dark, Sym_light: symbiotic anemones growing in 12h:12h light:dark photoperiod.

## Discussion

We documented the progress of oral regeneration in the symbiotic anemone Aiptasia (**Fig. 1; Fig. S2; S4**) and defined distinct stages that are more accurate in correlating the phenotype with the underlying cellular and molecular processes, than is relying solely on time after amputation (e.g., van der Burg et al., 2020). The steps and time to full oral regeneration were similar to those in the anemones studied by Singer & Palmer, 1969a, which took 7 to 10 days to regenerate. These authors also described a “pigmented ring” within 2 days after amputation, which they attributed to the migration of algae, and observed that tentacle buds arose from that ring (Singer & Palmer, 1969a). However, that change in the pigmentation or the distribution of the algae upon oral amputation was not evident in our observations.

Wound healing was reached within the first 8 to 24 hpa, regardless of symbiotic status and light regime; however, the time to tentacle bud formation was significantly shorter (ca. 12h difference) and less variable in symbiotic anemones under normal light than in aposymbiotic anemones or symbiotic anemones kept in the dark (**Fig. 2**). The most plausible explanation for this pattern is that the contribution to host nutrition by symbiont-derived photosynthates shortened regeneration time. The onset of tentacle budding significantly increases anemone metabolic demand, as measured by oxygen consumption (Singer & Palmer, 1969b), and likely requires cell proliferation (Singer, 1971), as supported by our observation that the addition of the mitotic inhibitor HU resulted in arrested oral regeneration of anemones at stage 3 (**Fig. S3**). In the temperate facultative symbiotic coral species *Astrangia poculata*, both nutrition and symbiotic state promoted healing and tissue recovery, although nutrition alone showed positive effects on coral recovery regardless of symbiotic state (Burmester et al., 2017, 2018). In our initial hypothesis, we proposed that signals from symbionts could promote regeneration by stimulating host cell proliferation (Tivey, Parkinson & Weis, 2020; Gorman et al., 2022).

However, our results suggest that symbiotic signals have little effect on the progress of oral regeneration and that feeding before amputation would abolish differences between symbiotic states.

Whole-body regeneration in Aipitasia occurs in the context of major injury (e.g., oral amputees) and also in asexual reproduction via pedal laceration. Pedal laceration has been used to generate clones in Aiptasia and to investigate the role of symbiosis in development (Presnell, Wirsching & Weis, 2022). There are distinct similarities and differences between the regeneration processes following oral amputation and pedal laceration. After healing the wound site, pedal lacerates undergo elongation and differentiated regeneration of mesenteries, column, oral disc, mouth, pharynx and tentacles (Presnell, Wirsching & Weis, 2022). Artificially generated pedal lacerates typically develop 10-12 tentacles by 14-21 days post-amputation compared to 7 days after oral amputation in adults. Those pedal lacerates also have a 30% chance of not regenerating in control conditions unlike in adults after oral amputation where nearly all animals regenerate (JB Cai, 2023, pers. comm.). The difference in regeneration rate and modality could be rooted in the type of wound and the total investment in regrowing a new polyp. In pedal laceration, the artificial wound is minor but the investment in regeneration is higher than in oral amputation, where a major wound is inflicted but the body column is retained. The commonalities and differences in both processes may help elucidate the role of algal symbionts in different homeostatic pathways.

## Conclusions

Aiptasia displays characteristic wound healing and high regenerative capacities found in other cnidarians. Here, we described distinct stages post-amputation from wound healing to oral regeneration within a week. Our results showed that photosynthate-derived nutrition from symbionts accelerated regeneration, but the effect was limited to only the onset of tentacle budding. These results suggest that oral regeneration in Aiptasia is primarily host-driven. Our study provides a reference for future research on the molecular mechanisms involved in Aiptasia oral regeneration by transcriptomic or proteomic analysis. Future comparative studies on different symbiotic cnidarians or different life stages could provide better insight into the role of symbiosis in regenerative processes.

## Declarations

## Acknowledgments

The authors wish to acknowledge the Confocal Microscopy Facility of the Center Quantitative Life Sciences at Oregon State University. We would like to thank animal care staff Alex Bridge, Lik Rong Lim, Olivia Arvas, Sarai Lopez and Daphnie Stacey for caring for the animals used in these experiments. We thank undergraduate researchers Kali Sivula, Riley Jones, Leah Thomas and Candice Vo, for their help in collecting data. We also thank Weis lab members, Erick White, Olivia Burleigh, Dr. Maria Ruggieri, Dr. Shumpei Maruyama and Dr. Valeri Sawiccy for their support and feedback.

## Authorship declaration

JBC, VMW and LP conceived and designed the experiments. JBC, SB and LP performed the experiments and analyzed the data. JBC and LP wrote the first draft of the manuscript and all authors commented on previous versions of the manuscript. All authors read and approved revised the final manuscript.

### Competing Interest statement

The authors declare that they have no competing interests.

### Funding statement

LP received support from “la Caixa” Foundation (ID 10010434), co-financed by the European Union’s Horizon 2020 research and innovation program under the Marie Sklodowska-Curie grant agreement No 847648, fellowship code is 104855. LP is funded by the Ramón y Cajal contract RYC2022-036761-I by MICIU/AEI/10.13039/5011 00011033 and the ESF+.

### Data availability

The raw data from all experiments is available in separate sheets in supplemental Data S1.

## Notes

### Competing Interest Statement

The authors have declared no competing interest.

