## Supplementary Material for "Aiptasia oral regeneration is host-controlled but supported by symbiont-derived photosynthates"

Corresponding Author:

### Supplementary Video

**Supplementary Video S1. Restoration of hydrostatic pressure upon oral amputation.** Time lapse during the fourth hour post oral amputation. Pictures were taken every 20s and then converted into a video at 20 frames per second.

### Supplementary Figures

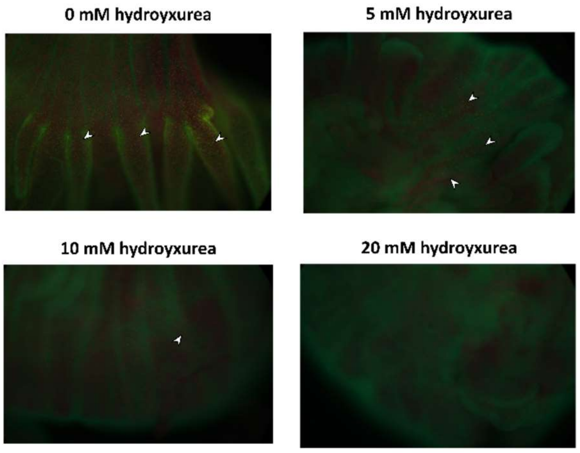


**Fig. S1.** Representative images of symbiotic animals from the hydroxyurea dosage tests viewed under the Olympus Vanox-T epifluorescence microscope. EdU-labelled cells (yellow with white arrowhead); host nuclei stained with Hoechst (green) and the autofluorescence from chlorophyll in algal symbionts (red).

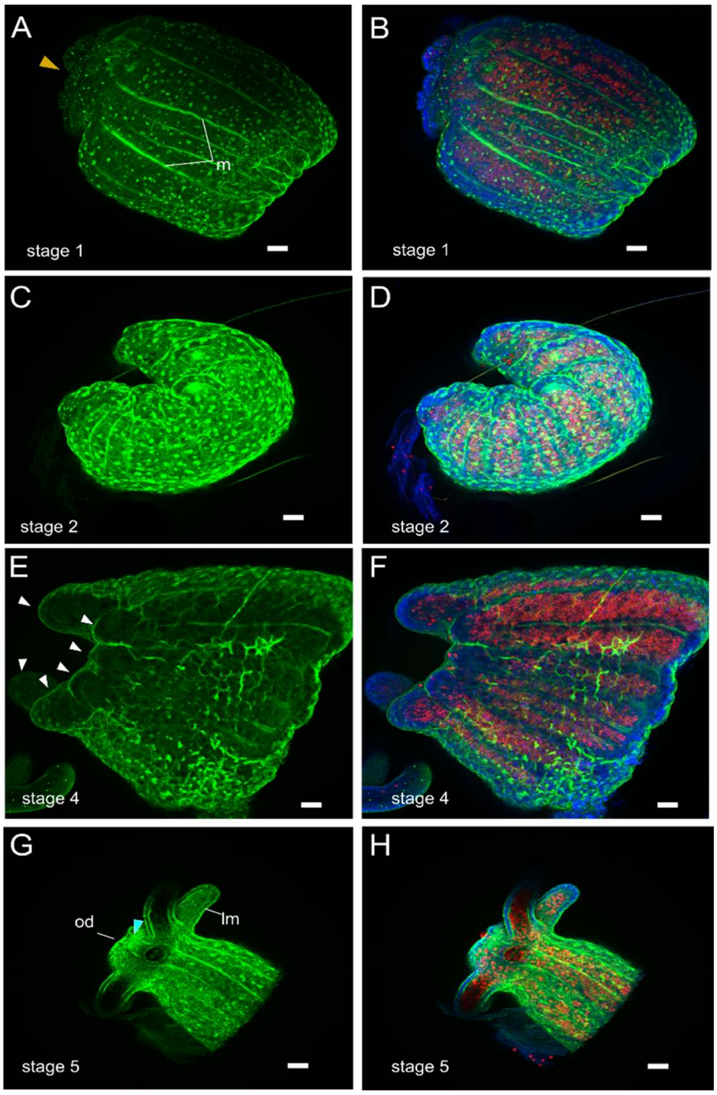


**Fig. S2. Regeneration of oral structures in Aiptasia.** At different stages of healing/regeneration, samples were fixed and processed for fluorescence staining of host cell nuclei (0.2 µg/mL Hoechst, Invitrogen) and actin filaments (0.0066 µM Alexa Fluor 488-phalloidin, Invitrogen) as in Presnell, Wirsching & Weis, (2022). (**A**) F-actin signal in anemones at stage 1, with open wound (yellow arrowhead). m: retractor muscle. (**B**) F-actin signal at stage 2. Epidermis rolled inward. (**C**) and (**D**) Stage 4 anemones showed the appearance of tentacle buds (arrowheads). (**E**) and (**F**) At stage 5, the tentacles had extended and their longitudinal musculature (lm) was formed. The oral disc (od) has regenerated and connected with the musculature in the body column (blue arrowhead). (**D**) and (**F**) show the superposing signals of F-actin, nuclei and algal symbionts.  Confocal z-stack projections. F-actin was stained with phalloidin (green) to visualize the cytoskeleton, host nuclei stained with Hoechst (blue) and algal symbionts are visible by chlorophyll autofluorescence (red). Scale bar: 50 µm.

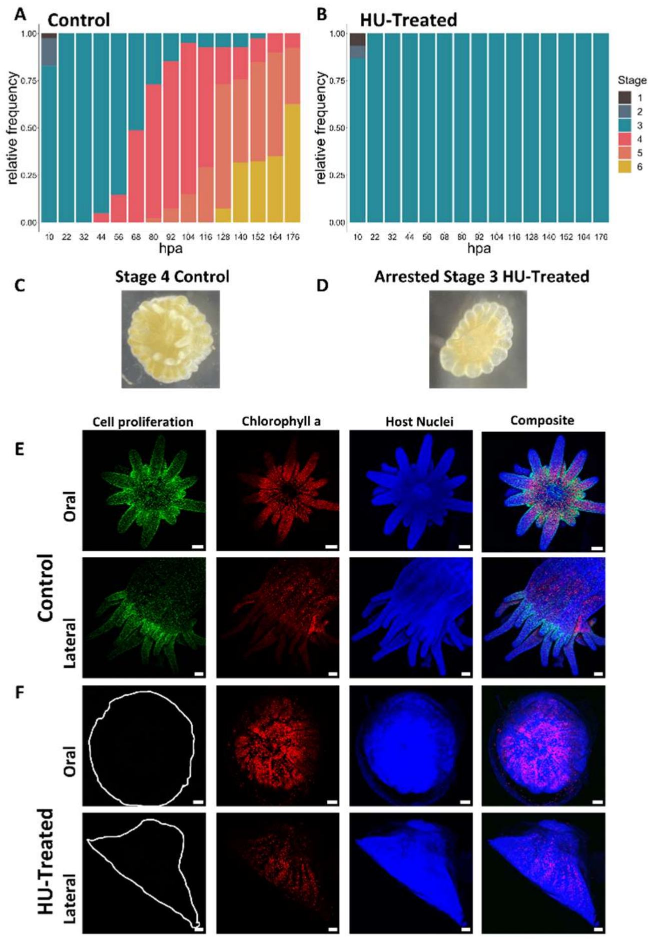


**Fig. S3. Cell division inhibitor treatment arrests regeneration.** Progress of regeneration over time in (A) the absence (0mM) or (B) presence (15mM) of mitotic inhibitor hydroxyurea (HU). Representative images of anemones in (C) control and (D) HU treatment at 80 hpa. (E) Control anemones and (F) HU-treated anemones at 176 hpa, after 24 h of EdU labeling. No EdU signal was detected in HU-treated anemones, polyp is outlined for reference. EdU-labelled cells (green); host nuclei stained with Hoechst (blue) and the autofluorescence from chlorophyll in algal symbionts (red). Scale bars: 100 µm.

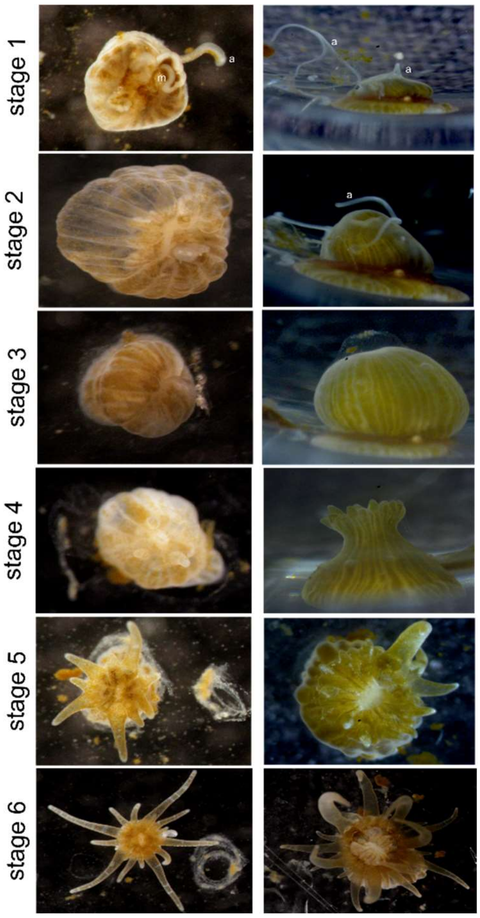


**Fig. S4. Representative images of the regeneration stages.** Stage 1: open wound (oral view), the acontia (a) and mesenteric filaments (m), and the retracted body column (lateral view). Stage 2: the wound is still open but epidermis is rolled inward (oral view) but the expansion of the body column suggests that the hydrostatic pressure was already restored. Stage 3: wound apparently closed and body column in the characteristic "onion shape". Stage 4: Tentacle buds appeared and the body column was elongated. Stage 5: the tentacles had outgrown and the regenerated oral disc is visible. Stage 6: same number of tentacles as before amputation.

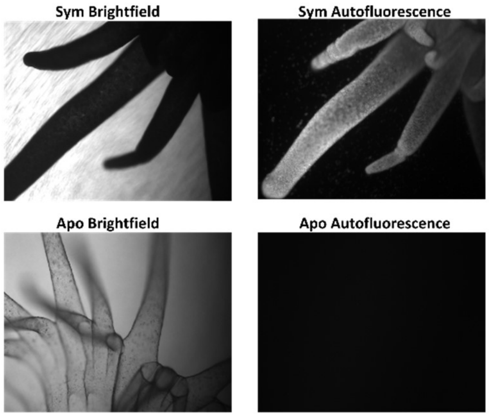


**Fig. S5. Representative images of symbiotic animals and aposymbiotic animals.** Polyps viewed under a Zeiss Axio Observer A1 fluorescence microscope with a black-and-white brightfield view and a view of the chlorophyll autofluorescence channel.

**Supplementary Table**

**Supplementary Table S1. Average time to reach stages 3, 4, 5, and 6 for each treatment.** For each stage, homoscedasticity of variances was tested by Levene's test. When test was non-significant (p-value ≥0.05), statistical comparison of time to reach stage was performed by non-parametric Kruskal-Wallis Rank sum test (KW). For stages 4 and 5, Levene's test reported heteroscedasticity; thus, we applied a non-binomial glm test; followed by Tukey post-hoc test, if significant (stage 4). Letters represent grouping according to Tukey's test results. df: degrees of freedom, P: p-value. Apo: aposymbiotic anemones; Sym_dark: symbiotic anemones growing in the dark, Sym_light: symbiotic anemones growing in 12h:12h light:dark photoperiod.

| Stage | Treatment | Mean±SD (hpa) | Median (hpa) | n | Statistical test |
|---|---|---|---|---|---|
| 3 | Apo | 10.7 ± 8.8 | 8 | 55 | KW Chi-squared = 2.2905; df = 2; P = 0.3181 |
| | Sym_dark | 13.4 ± 8.8 | 8 | 23 | |
| | Sym_light | 9.6 ± 6.8 | 8 | 47 | |
| 4 | | | | | df = 2, P = **2.148e-04** |
| | Apo | 43.2 ± 12.5 | 48 | 48 | **(a)** |
| | Sym_dark | 45.5 ± 15.1 | 48 | 22 | **(a)** |
| | Sym_light | 34.8 ± 9.3 | 32 | 40 | **(b) p < 0.005** |
| 5 | Apo | 90.6 ±33.7 | 84 | 40 | df = 2; P = 0.1406 |
| | Sym_dark | 77.6 ± 13.5 | 72 | 17 | |
| | Sym_light | 81.6 ± 27.3 | 72 | 41 | |
| 6 | Apo | 140.3 ± 31.6 | 168 | 46 | KW Chi-squared = 2.299, df = 2, P = 0.3168 |
| | Sym_dark | 150.7 ± 29.5 | 168 | 18 | |
| | Sym_light | 136.9 ± 31.9 | 120 | 37 | |
